# Calcitriol and Non-Calcemic Vitamin D Analogue, 22-Oxacalcitriol, Attenuate Developmental and Pathological Ocular Angiogenesis Ex Vivo and In Vivo

**DOI:** 10.1101/515387

**Authors:** SL Merrigan, B Park, Z Ali, LD Jensen, TW Corson, BN Kennedy

## Abstract

Aberrant ocular blood vessel growth can underpin vision loss in leading causes of blindness, including neovascular age-related macular degeneration, retinopathy of prematurity and proliferative diabetic retinopathy. Current pharmacological interventions require repeated invasive administrations, lack efficacy in some patients and are associated with poor patient compliance and tachyphylaxis. Small molecule vitamin D has de novo pro-differentiative, anti-proliferative, immunomodulatory, pro-apoptotic and anti-angiogenic properties. Here, our aim was to validate the anti-angiogenic activity of the biologically active form of vitamin D, calcitriol, and a selected vitamin D analogue, 22-oxacalcitriol, across a range of ocular angiogenesis models.

First, we validated the anti-angiogenic activity of calcitriol, showing calcitriol to significantly inhibit choroidal sprouting in an *ex vivo* mouse choroidal fragment sprouting assay. Viability studies in human RPE cell line, ARPE-19, suggested non-calcemic vitamin D analogues have the least off-target anti-proliferative activity compared to calcitriol and additional analogues. Thereafter, the ocular anti-angiogenic activity of non-calcemic vitamin D analogue, 22-oxacalcitriol, was demonstrated in the *ex vivo* mouse choroidal fragment sprouting assay. In zebrafish larvae, 22-oxacalcitriol was anti-angiogenic, inducing a dose-dependent reduction in choriocapillary angiogenesis. Inhibition of mouse retinal vasculature development was not induced by systemically delivered calcitriol. However, both calcitriol and 22-oxacalcitriol administered intraperitoneally significantly attenuate choroidal neovascularisation lesion volume in the laser-induced CNV mouse model. 22-oxacalcitriol presented with a more favourable safety profile than calcitriol.

In summary, calcitriol and 22-oxacalcitriol attenuate *ex vivo* and *in vivo* choroidal vasculature angiogenesis. Vitamin D has potential as a preventative or interventional treatment for ophthalmic neovascular indications.

## Introduction

Growth of pathological ocular blood vessels can underpin vision loss in leading causes of blindness including neovascular age-related macular degeneration (nAMD) and proliferative diabetic retinopathy (PDR). Worldwide 8.7% of blindness results from AMD. nAMD accounts for 10% of AMD cases but >80% of cases with poor visual acuity [1-4]. nAMD and rapid vision loss is driven by pathological choroidal vasculature angiogenesis. This pathological choroidal vasculature can be deficient in tight junctions, leak plasma or blood, cause scarring, project through the Bruch’s membrane, cause retinal pigmented epithelium (RPE) detachment and disrupt normal perfusion of the retina [5-7]. Worldwide 382 million people suffer from diabetes, ∼35% of whom develop DR, making this the leading cause of blindness in the working age population [8]. Severe vision loss is a consequence of macular oedema and the sprouting of poorly formed retinal vasculature into the vitreous [8]. Retinal neovascularisation triggered by insufficient perfusion can result in haemorrhaging and retinal detachment [9].

Endogenous pro-angiogenic factors typified by vascular endothelial growth factor (VEGF), angiopoietin (ang-1, ang-2), platelet-derived growth factor (PDGF-A, PDGF-B) and transforming growth factor (TGF-β), plus their cognate receptors, promote normal vasculature development [6]. After development, a tightly regulated balance of pro- and anti-angiogenic factors maintain the mature quiescent vasculature. Pathological insults such as hypoxia can disrupt this equilibrium and promote neovascularisation [10, 11]. VEGF is a pivotal regulator of nAMD, clinically evident from the success of anti-VEGF targeting therapies. Ranibizumab (LUCENTIS^®^), bevacizumab (Avastin^®^) and aflibercept (Eylea^®^) are utilised in the treatment of ocular neovascularisation [12]. These anti-VEGF therapies cause vessel regression and improve visual function [4]. Despite representing the standard of care, several treatment limitations exist. Firstly, with molecular weights between 50-149 kDa, current interventions require administration via intravitreal injection [13]. This places a burden both on patients and clinicians, resulting in inadequate dosing, exemplified by the CATT/IVAN trial, where an average of 4-5 treatments were administered compared to the recommended 7-8 [7]. Secondly, repeat administrations are required: aflibercept has the greatest intravitreal half-life yet injections are required every 2 months [4, 14]. Thirdly, anti-VEGF therapies are associated with a severe economic burden, with aflibercept costing approximately €1,000 per injection [15]. Finally, anti-VEGF therapy can lack efficacy in “non-responsive” populations and tachyphylaxis can occur. A non-responsive population of 45% is reported for bevacizumab [16]. Tachyphylaxis is postulated to be a consequence of compensatory VEGF upregulation or generation of neutralising antibodies [14]. These limitations highlight the need to identify and develop safe, efficacious, cost effective anti-angiogenics with a less invasive route of administration.

Vitamin D is a fat-soluble vitamin and steroid hormone with pleiotropic health implications. Recognised as having pro-differentiative, anti-proliferative, immunomodulatory, pro-apoptotic and anti-angiogenic properties; vitamin D is under examination for malignant, cardiovascular, cognitive, metabolic, infectious and autoimmune indications [17-19]. Interestingly, the vitamin D receptor (VDR) is expressed in the cornea, lens, ciliary body, RPE, ganglion cell layer and photoreceptors, supporting ocular functions [20]. In 2017, we reported calcitriol (vitamin D) and diverse VDR agonists including vitamin D_2_ analogues, vitamin D_3_ analogues and a pro-hormone to attenuate *in vivo* ocular vasculature development in zebrafish [21]. Further interrogation of the anti-angiogenic activity was needed in mammalian models to assess the therapeutic potential of vitamin D.

Here, we examined the anti-angiogenic activity of calcitriol or 22-oxacalcitriol in choroidal or retinal vasculature systems, *in vitro, ex vivo* and *in vivo*. Calcitriol and 22-oxacalcitriol significantly inhibit *ex vivo* mouse choroidal sprouting angiogenesis, yet in a simpler, non-ocular *in vitro* tubule formation assay anti-angiogenic activity was not identified. With increased model complexity calcitriol and/or 22-oxacalcitriol again induced anti-angiogenic responses, showing reduced angiogenesis in an *in vivo* zebrafish model and attenuated neovascularisation in an *in vivo* mouse model that recapitulates features of neovascular age-related macular degeneration, the laser-induced choroidal neovascularisation model (L-CNV). Drug safety was assessed through animal weight monitoring and 22-oxacalcitriol presented with a superior safety profile compared to calcitriol. Our comprehensive studies presented here support further pre-clinical investigations into non-calcaemic vitamin D analogue, 22-oxacalcitriol, for the treatment or prevention of choroidal neovascularisation.

## Methods

### Mouse choroidal sprouting angiogenesis assay

The mouse choroidal sprouting procedure was adapted from Shao et al [22]. C57BL/6J mice aged between 6-12 weeks were euthanised by CO_2_ asphyxiation, eyes immediately enucleated and placed in ice cold Endothelial Cell Growth Medium (PromoCell). Eyes were cut along the pars planar, cornea removed, lens removed, and four incisions made facilitating flattening of the eyecup. The neural retina was removed from the RPE/choroid complex and six-eight 1 x 1 mm RPE-choroid explants cut from each quadrant. Explants were transferred to 30 µl thawed Matrigel^®^ (BD Biosciences) in a 24 well plate, uniformly orientated, incubated for 20 min at 37°C (5% CO_2_, 95% O_2_) and 500 μl medium applied. Following 1 day of culturing, medium was exchanged, and vehicle control, calcitriol and 22-oxacalcitriol treatments applied with a final well volume of 500 μl. Treatments were replenished on day 3-4. Culturing was ended on day 7 and explants were imaged live, calcein stained or fixed in 4% PFA overnight.

### Calcein staining

Following 7 days of culturing, RPE-choroid explants were washed with phosphate-buffered saline (PBS) and 300 μl of 8 µg/ml Calcein AM (Thermo Fisher Scientific) applied to each explant and incubated for 1 h at 37°C (5% CO_2_, 95% O_2_). Three PBS washes were performed, and explants imaged.

### Image acquisition and sprouting area quantification

Brightfield images were acquired using Olympus SZX16 or Zeiss Axiovert 200 M microscopes with Cell^F or Zeiss Axiovision image analysis software. Sprouting area and explant area were manually quantified using ImageJ freehand tool and explant area subtracted from overall area. Statistical differences between vehicle- and drug-treated samples were determined by one-way ANOVA with Dunnett’s post-hoc test. Statistical analyses were performed with PRISM 5 software and significance accepted where P≤0.05.

### ARPE-19 cell MTT cell viability assay

ARPE-19 cells were maintained in Dulbecco’s Modified Eagle’s Medium Nutrient Mixture F-12 (Sigma Aldrich) with 10% fetal bovine serum (FBS) (Gibco), 2 mM L-glutamine (Gibco) and 100 units/mL penicillin-streptomycin (Gibco). Briefly, 1.4 × 10^4^ cells were applied per well of a 96 well plate and incubated at 37°C (5% CO_2_, 95% O_2_) overnight in compete medium. Complete medium was replaced with FBS-negative medium and plates incubated overnight at 37°C (5% CO_2_, 95% O_2_). FBS-negative medium was replaced with calcitriol treatments in FBS-negative medium at indicated concentrations, with 4 replicate wells per treatment and incubated at 37°C (5% CO_2_, 95% O_2_). At the desired treatment end-point 10 µl of 5 mg/ml MTT labelling Reagent (Roche) was applied per well and plates incubated at 37°C (5% CO_2_, 95% O_2_) for 2 h. 100 µl solubilisation solution (Roche) was applied per well and plates incubated at 37°C (5% CO_2_, 95% O_2_) for 4 h and absorbance read at 570 nm (microplate reader, mtx lab systems). Statistical differences between vehicle- and drug-treated samples were determined by one-way ANOVA with Dunnett’s post-hoc test. Statistical analyses were performed with PRISM 5 software and significance accepted where P≤0.05.

### Choriocapillaris development assay in zebrafish

Fertilised Tg(*Fli1a:EGFP*)^y1^ transgenic zebrafish eggs were incubated in Phenylthiourea -containing E3-medium for 24 hours and then treated with 0.1% DMSO (vehicle control) or 22-oxacalcitriol (Cayman Chemicals) at 0.1, 1.0 or 10 µM for an additional 48 hours. The larvae were then anesthetised with 0.04% MS-222 (Ethyl 3-aminobenzoate methane sulfonic acid salt 98%, Sigma Aldrich) and fixed in 4% PFA (Sigma Aldrich) for 30 minutes at room temperature. The eyes were removed and dissected using watchmakers’ forceps (Dumont #5) under a dissection microscope (Nikon SMZ 1500). The eyes were flat mounted on glass slides in Vectashield mounting medium (H-1000 Vector laboratories) and imaged by confocal microscopy (Zeiss, LSM 700). The number of interstitial pillars (ISPs) were counted manually and used as a measure of the extent of active vascular growth.

### Calcitriol tolerance study in adult mice: Weight and Retina histology

Calcitriol (Selleckchem) was dissolved to 1 mg/ml in ethanol and working dilutions to 1 µg/ml in PBS prepared. Male C57BL/6J mice between 3-6 months were subcutaneously (s.c.) administered 50 ng calcitriol or vehicle control and animal weight recorded daily over a 7 day period.

On day 7, mice were euthanised by CO_2_ asphyxiation, eyes enucleated, bisected and fixed with 2.5%: 2% glutaraldehyde: PFA in 0.1 M Sorenson phosphate buffer (SPB) (pH 7.3). Fixed eyes were washed in 0.1 M SPB and transferred to 1% osmium tetroxide in 0.2 M SPB for 1 h at room temperature. Eyes were exposed to an ethanol gradient; 30%, 50%, 70%, 90% for 10 min each; 100% for 60 min and acetone for 30 min. Eyes were uniformly positioned within an Epon resin primed mould, mould levelled with Epon resin composed of agar 100 resin, dodecenyl succinic anhydride, methyl nadic anhydride and 2,4,6-tris (dimethylaminomethyl) (Agar Scientific) and incubated overnight at 50 °C. Ultra-thin ocular cross sections, 1 µm, were acquired using a diamond knife and Leica EM UC6 microtome. Sections were toluidine blue stained, cover-slipped (DPX mounting medium) and representative images acquired (Nikon Eclipse E80i Microscope, Canon camera). Mouse retina morphology between vehicle control and calcitriol treated samples was compared to identify deviations in retina cell organisation, retinal thickness and pyknotic nuclei presence.

### Mouse model of retinal vasculature development

Experimental design followed Yagasaki et al [23]. Dams along with their pups were raised in standard light (12 h light and 12 h dark cycle) and standard air conditions for the duration of the study.

#### Calcitriol dosing

Calcitriol was prepared as previously reported [24]. Calcitriol (Selleckchem) was dissolved to 1 mg/ml in ethanol and working dilutions to 1 µg/ml in PBS prepared. Calcitriol multiple injection study: Pups received a 3.75 ng calcitriol or vehicle control s.c. treatment on P1, P3, P5 and P7. Animal welfare was monitored daily until P4 or P8. On P4 and P8 mouse pups were euthanised by cervical dislocation, eyes immediately enucleated and fixed with 4% PFA overnight at 4°C.

#### Retina flatmount

Fixed eyes were positioned on a made-for-purpose indented dental wax strip, in PBS under a dissecting microscope. The eye was gripped at the optic nerve with a Dumont no. 5 forceps and excess exterior muscle removed using a springbow microdissection scissors. The optic nerve was removed, the eye was pierced along the pars planar, the anterior eye removed using a springbow dissection scissors and lens removed. The remaining eyecup was transferred to a glass slide and 4 incisions made 2 mm from the site of the optic nerve with a no. 11 scalpel blade dividing the retina into 4 quadrants. Using a no. 11 blade the periphery of the quadrants was cut, straightening the outer edge and preventing curling of the retina. Retinal flat mounts were stored in perm/block buffer composed of PBS with 0.5% Triton-X100, 1% goat serum, and 0.1 mM CaCl_2_ (Sigma-Aldrich).

#### Isolectin staining

Isolectin staining was performed as previously reported [25]. Retina flat mounts underwent permeabilization with perm/block buffer overnight at 4°C. Perm/block buffer was replaced with 20 µg/ml GS isolectin B4 (ThermoFisher Scientific) in perm/block buffer and incubated overnight at 4°C. Flat-mounts underwent 8 perm/block buffer washes over 4 hours at 37°C with 30 min interval changes. Flat mounts were stained with Alexa-streptavidin-564 (Thermo Fisher Scientific) diluted in perm/block buffer 1:500 overnight at 4°C. Flat-mounts underwent 8 perm/block buffer washes over 4 hours at 37°C with 30 min interval changes. Flat mounts were stored in PBS with 0.1 mM CaCl_2_. Flat-mounts were transferred to a glass slide, the retina cover-slipped with Aqua-Poly mount (Polyscience Inc) and stored protected from light at 4°C.

#### Flat-mount image acquisition and retinal vasculature area quantification

Fluorescent images were acquired using a Zeiss AxioVert 200M fluorescent microscope, Andor IQ2 software with Andor montaging or Olympus SZX16 fluorescence microscope with Cell^F software. Retinal superficial vasculature development was expressed as vasculature area compared to total flat mount area. Area measurements were performed using ImageJ software freehand tool.

### Mouse Model of Laser-Induced Choroidal Neovascularisation

Wild-type female C57BL/6J mice, 6–8 weeks of age, were purchased from the Jackson Laboratory (Bar Harbor, ME), and housed under standard conditions [26]. Intraperitoneal injections (i.p.) of 60 mg/kg ketamine hydrochloride and 2.5 mg/kg xylazine mixture were used for anaesthesia, and isoflurane overdose for euthanasia. Body weights were determined daily. All analyses were performed by a masked investigator.

#### Calcitriol dosing

Calcitriol (1000 ng/mL in almond oil; Professional Compounding Centers of America, Houston, TX) was purchased from Indiana School of Medicine Laboratory Animal Resource Center’s drug distribution center. Each mouse received once-daily i.p. injections of 5 µg/kg calcitriol or almond oil vehicle for 14 days (5 days on/2 days off). The dose was determined based on published observations [27].

#### 22-oxacalcitriol dosing

Pure crystalline solid 22-oxacalcitriol (Cayman Chemical, Ann Arbor, MI) was dissolved in ethanol as previously reported to yield a 2 µg/µL stock solution [28, 29]. This stock solution was diluted in PBS to working solution, 2 µg/mL 22-oxacalcitriol in 0.1% ethanol-PBS on day of the injection. Each mouse in the treatment group received once-daily i.p. injections of 15 µg/kg 22-oxacalcitriol or ethanol-PBS vehicle every day for 14 days.

#### Laser-induced choroidal neovascularisation

The L-CNV mouse model was performed as previously described [30]. Briefly, both eyes of 6-8 week old C57BL/6J mice were dilated using tropicamide, and subjected to laser treatment using 50 µm spot size, 50 ms duration and 250 mV pulses of an ophthalmic argon green laser wavelength 532 nm, coupled to a slit lamp. Three laser burns per eye were created around the optic nerve at 12, 3 and 9 o’ clock positions. Optical coherence tomography (OCT) was performed in L-CNV mice as described previously [30], on days 7 and 14 post laser, using a Micron III intraocular imaging system (Phoenix Research Labs, Pleasanton, CA, USA). Briefly, eyes of anesthetised mice were dilated with 1% tropicamide solution (Alcon, Fort Worth, TX, USA) and lubricated with Gonak hypromellose ophthalmic solution (Akorn, Lake Forest, IL, USA). Horizontal and vertical OCT images were taken per lesion and L-CNV lesion volumes were obtained using the quantification method previously established [30, 31]. To assess vascular leakage, fluorescein angiography was performed on day 14 post L-CNV by i.p. injection of 50 μL of 25% fluorescein sodium (Fisher Scientific, Pittsburgh, PA, USA). Fundus images were taken using the Micron III system and Streampix software.

#### Choroidal flatmount

Mouse eyes were harvested on day 14 post L-CNV induction. The eyes were enucleated and fixed in 4% PFA in PBS for an hour at 4°C. The anterior portion including lens and the retina were removed, then the posterior eyecups were dissected out and underwent further fixation in 4% PFA in PBS overnight. The fixed eye cups were washed in blocking buffer (0.3% Triton X-100, 5% bovine serum albumin (BSA) in PBS) for two hours at 4°C. The eye cups were then stained for vasculature using the rhodamine-labeled *Ricinus communis* agglutinin I (Vector Labs, Burlingame, CA, USA) and Alexa Fluor 488 conjugated-Isolectin B4 from *Griffonia simplicifolia* (GS-IB4) (Molecular Probes, Thermo Fisher Scientific) at 1:250 dilution in buffer containing 0.3% Triton X-100, 0.5% BSA in PBS, overnight at 4°C. The posterior eyecups were then washed three times with PBS and mounted in fluorescent mounting medium (VectaShield; Vector Laboratories, Inc.) and cover-slipped. Confocal imaging and analysis of L-CNV lesion volume were performed as previously described [31]. Treatments were compared by unpaired T-test (two tailed) with Welch’s correction, while mouse body weights were compared by two-way repeated-measures ANOVA with Holm-Sidak post hoc tests using GraphPad Prism v. 6.

### Mouse eye RNA extraction, cDNA synthesis and QRT-PCR

Mouse eyes were harvested on day 14 post L-CNV induction, lens and cornea removed and eye stored in RNAlater. Total ocular RNA was extracted using mirVana™ miRNA Isolation Kit (ThermoFisher Scientific) as per the manufacturer’s instructions. cDNA synthesis was carried out using VILO cDNA Synthesis Kit (Thermo Fisher Scientific) as per the manufacturer’s instructions. QRT-PCR reactions: 0.5 μL Taqman specific probe, 5 μL TaqMan Gene Expression Master Mix, 2.5 μL RNAse-free water and 2 μL cDNA template were made up on ice. QRT-PCR cycles were carried out with a QuantStudio 7 Flex Real-Time PCR System with QuantStudio™ Software and the following conditions applied: 50°C for 2 min, 95°C for 10 min, 95°C for 15 s with 40 repeats and 60°C for 1 min.

## Results

### Calcitriol attenuates mouse *ex vivo* choroid-RPE fragment sprouting angiogenesis

Previously, we demonstrated calcitriol and seven other VDR agonists to inhibit ocular vasculature development in zebrafish larvae [21]. To identify the most active anti-angiogenic VDR agonist in mammalian models, the tubule formation assay, a late stage *in vitro* angiogenesis model, was performed. Human dermal-derived microvascular endothelial cells, HMEC-1 cells, were seeded in matrix and cultured with 10 µM calcitriol, 22-oxacalcitriol, tacalcitol or vehicle control and tubule formation quantified after 16 h. The Angiogenesis Analyzer for ImageJ was utilised for automatic unbiased measurement of tubule formation properties. Surprisingly, VDR agonist-treated HMEC-1 cells exhibited no significant difference in tubule formation compared to vehicle controls **(Supp Figure 1A-B)**. Tubule formation properties are influenced by cell type (primary or immortalised), derivation (human or non-human) and tissue origin [32]. With ocular selective anti-angiogenic activity previously identified in zebrafish larvae, tubule formation was also investigated in human retinal-derived microvascular endothelial cells (HREC). HREC cells were seeded in a matrix and cultured with 10 µM calcitriol for 16 h. Again, no significant tubule formation difference was identified between vehicle control and calcitriol treated HREC cells **(Supp Figure 1C-D**).

**Figure 1:**
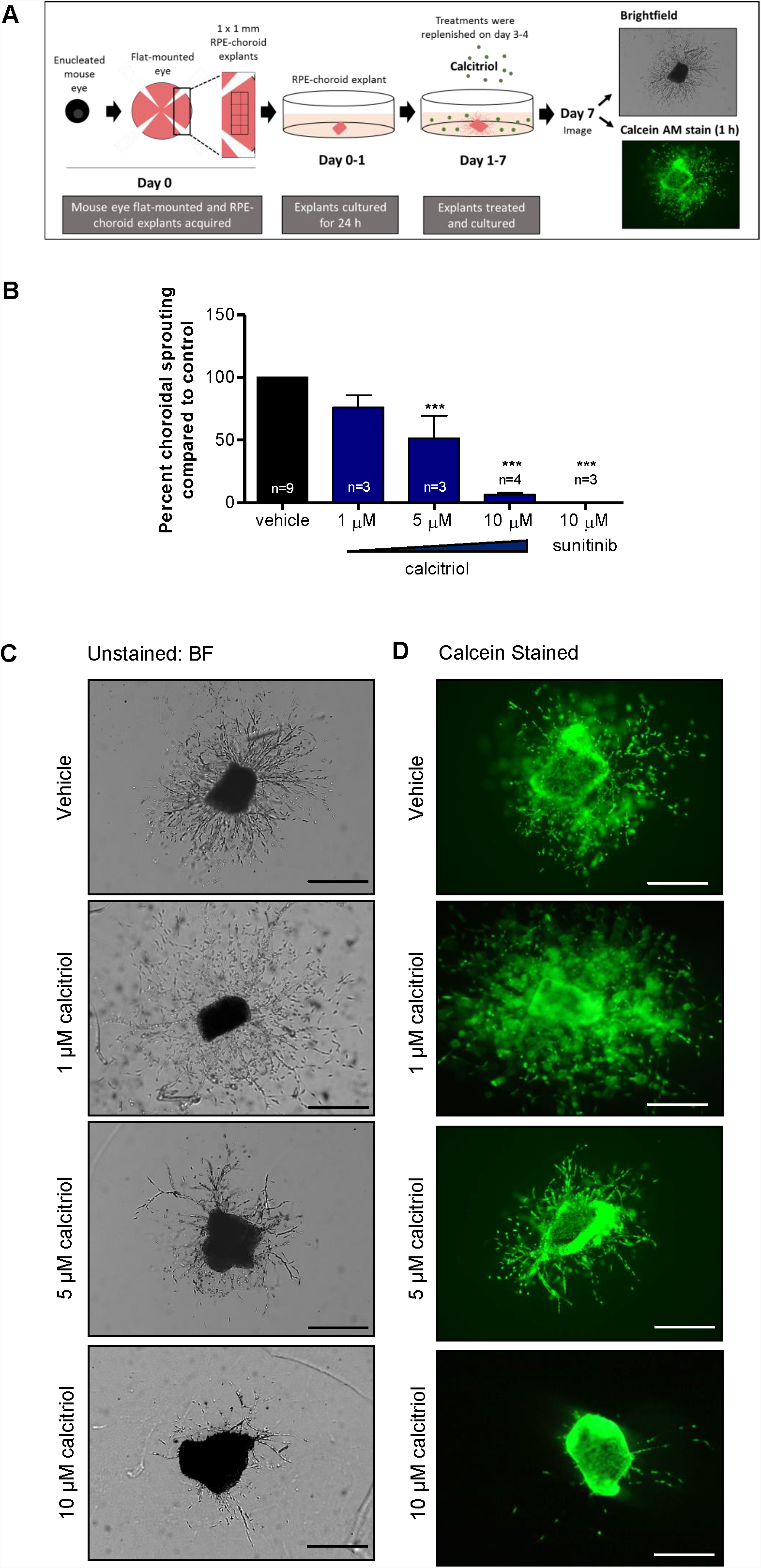
Calcitriol attenuates mouse choroidal sprouting angiogenesis. **(A)** Mouse RPE-choroidal fragments were cultured in Matrigel^®^ for 24 h and further cultured with indicated drug treatments for 6 days. On day 7, samples were fixed and sprouting area quantified from phase contrast images using ImageJ freehand tool. **(B)** Calcitriol 5-10 µM or 10 µM sunitinib positive control significantly attenuated choroidal sprouting angiogenesis. Graph shows mean percent sprouting area compared to vehicle control ±SEM; statistical analyses were performed using one-way ANOVA with Dunnett’s post hoc test, asterisk indicates ***p≤ 0.001 and n as indicated with up to 6 replicates per individual experiment. **(C)** Representative brightfield (BF) images of mouse choroidal sprouting after 7 days with indicated treatments. **(D)** Calcein stained representative images of mouse choroidal sprouting after 7 days with indicated treatments. Calcein staining of RPE-choroidal cultures confirms explant viability in vehicle and calcitriol treated explants. Scale bar represents 0.5 mm.

To investigate the anti-angiogenic activity of calcitriol in a more physiologically relevant model, the *ex vivo* mouse choroidal sprouting angiogenesis assay was employed. This system is multicellular in nature and accounts for micro-environmental cues which support angiogenesis [22]. Calcitriol treatments between 5-10 µM significantly (p<0.001) reduced choroidal sprouting area by up to 93% compared to vehicle control. No significant difference in sprouting was identified with 1 µM calcitriol treatments **(Figure 1B-D)**. Calcein staining confirmed explant and sprout viability after 1-10 µM treatments **(Figure 1D)**.

### Calcitriol attenuates RPE cell viability, while non-calcemic vitamin D_3_ analogues show a greater RPE cell safety profile

Pro-apoptotic and anti-proliferative properties of calcitriol are known [33]. Such actions on endothelial cells could underpin the anti-angiogenic mechanism of calcitriol. However, induction of apoptosis is undesirable in neighbouring cells such as the RPE. Thus, we sought to identify VDR agonists with negligible effects on RPE cell viability. VDR agonist-induced changes in ARPE-19 cell number were determined by the surrogate measure of metabolic activity, quantified using the MTT assay. Calcitriol was tolerated over 24 h in ARPE-19 cells, with no significant change in cell viability with concentrations ≤20 µM **(Figure 2A)**. However, treatments with ≥10 µM calcitriol for 48 h significantly reduced ARPE-19 cell viability in a concentration-dependent manner, 10 µM (p≤0.05), 15 µM (p≤0.01) and 20 µM (p≤0.001) **(Figure 2B)**. Cell viability in response to a range of VDR agonist treatments was subsequently investigated in ARPE-19 cells over 96 h. Vitamin D_2_ analogue, doxercalciferol, reduced ARPE-19 cell viability (∼42%) only with 10 µM treatment (p≤0.01) **(Figure 2C)**. Vitamin D_2_ analogue, paricalcitol, had no significant effect on APRE-19 cell viability with treatments ≤10 µM **(Figure 2D)**. Vitamin D_3_ analogues, tacalcitol and calcipotriol induced a significant reduction (∼42 and 29%, respectively) in ARPE-19 cell viability with concentrations ≥5 µM **(Figure 2E-F)**. Non-calcemic vitamin D_3_ analogues were better tolerated. No significant change in ARPE-19 cell viability was identified with 22-oxacalcitriol or EB 1089 treatments ≤10 µM **(Figure 2G-H)**. Therefore, non-calcemic vitamin D_3_ analogue 22-oxacalcitriol was selected for further investigation. Finally, vitamin D_3_ pro-hormone, calcifediol, significantly attenuated ARPE-19 cell viability by up to 45% with concentrations ≥5 µM (p≤0.001) **(Figure 2I)**.

**Figure 2:**
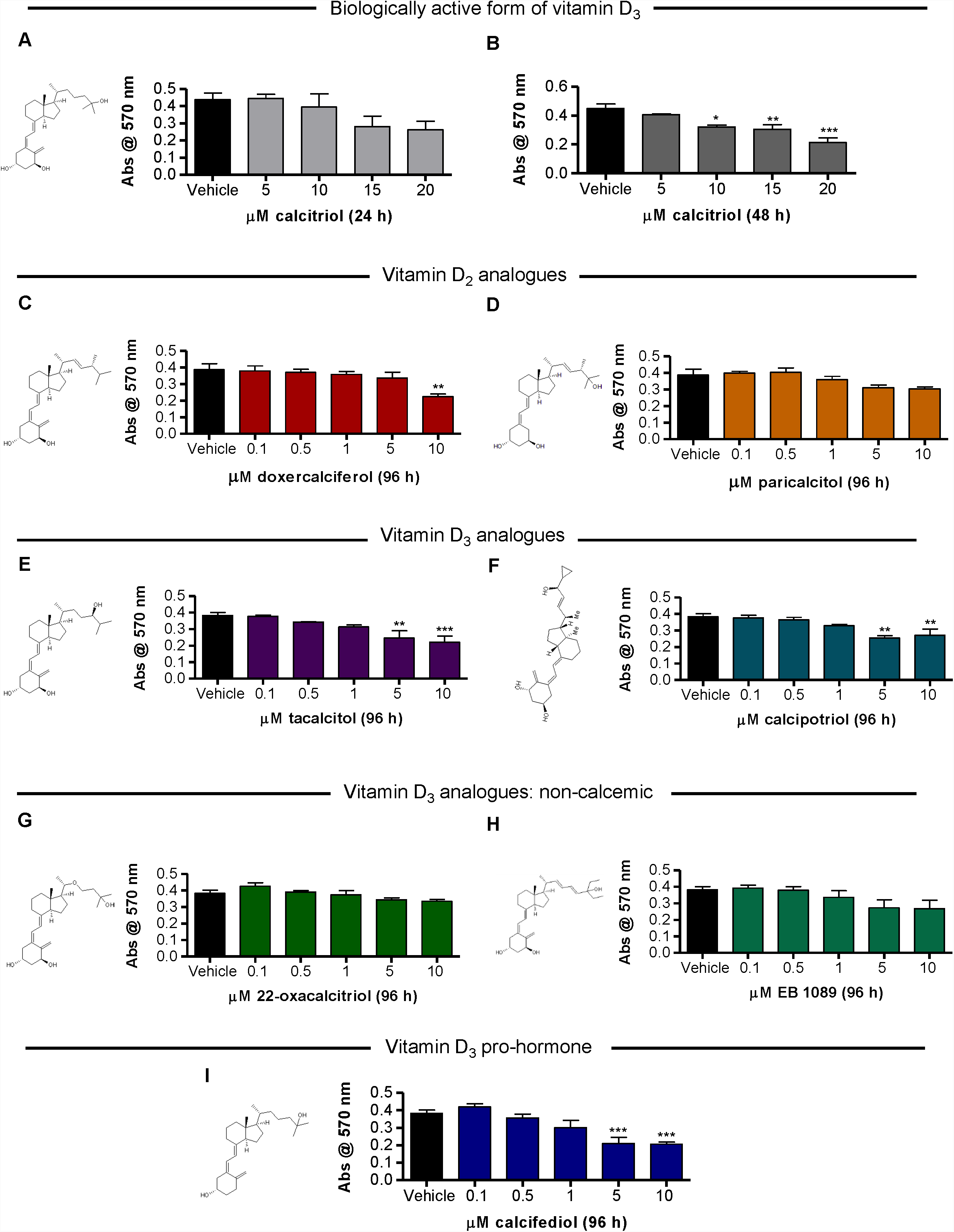
Vitamin D analogues attenuate ARPE-19 cell viability in a time- and concentration-dependent manner. **(A-B)** Serum starved ARPE-19 cells were cultured with 5-20 µM calcitriol for 24 h or 48 h and cell viability assessed by MTT assay. No significant difference in cell metabolic activity with 24 h calcitriol treatment was found. Calcitriol treatments (10-20 µM) for 48 h significantly reduced cell metabolic activity. **(C-I)** Serum starved ARPE-19 cells were cultured with 0.1-10 µM vitamin D_2_ analogues, vitamin D_3_ analogues, non-calcemic vitamin D_3_ analogues, vitamin D3 pro-hormone or vehicle control for 96 h and cell viability assessed by MTT assay. **(C-D)** Graphs show mean absorbance at 570 nm in response to vitamin D_2_ analogue treatment. **(C)** ARPE-19 cell viability was attenuated by 10 µM doxercalciferol treatment. **(D)** No significant change in ARPE-19 cell viability was identified with 0.1-10 µM paricalcitol treatment. **(E-F)** Graphs show mean absorbance at 570 nm in response to vitamin D_3_ analogue treatment. **(E)** ARPE-19 cell viability was attenuated by 5-10 µM tacalcitol treatment. **(F)** ARPE-19 cell viability was attenuated by 5-10 µM calcipotriol treatment. **(G-H)** Graphs show mean absorbance at 570 nm in response to non-calcemic vitamin D_3_ analogue treatment. No significant change in ARPE-19 cell viability was identified with 0.1-10 µM 22-oxacalcitriol or EB 1089 treatment. **(I)** Graphs show mean absorbance at 570 nm in response to vitamin D_3_ pro-hormone, calcifediol. ARPE-19 cell viability was attenuated by 5-10 µM calcifediol treatment. Graphs show mean absorbance at 570 nm ±SEM; statistical analysis by one-way ANOVA with Dunnett’s post-hoc test compared to vehicle control; asterisk signifies *p≤0.05, ** p≤0.01 and *** p≤0.001 and group size is n=3, with N=4/n.

### Non-calcemic Vitamin D_3_ analogue 22-oxacalcitriol inhibits mouse *ex vivo* choroidal sprouting angiogenesis

To investigate the anti-angiogenic activity of non-calcemic Vitamin D_3_ analogue 22-oxacalcitriol, the *ex vivo* choroidal sprouting angiogenesis assay was again performed. 10 µM 22-oxacalcitriol significantly attenuated mouse *ex vivo* choroid-RPE fragment sprouting angiogenesis by up to 42 % (p≤0.05) **(Figure 3A-C)**. In addition, calcein staining confirmed explant and sprout viability after treatment **(Figure 3C)**.

**Figure 3:**
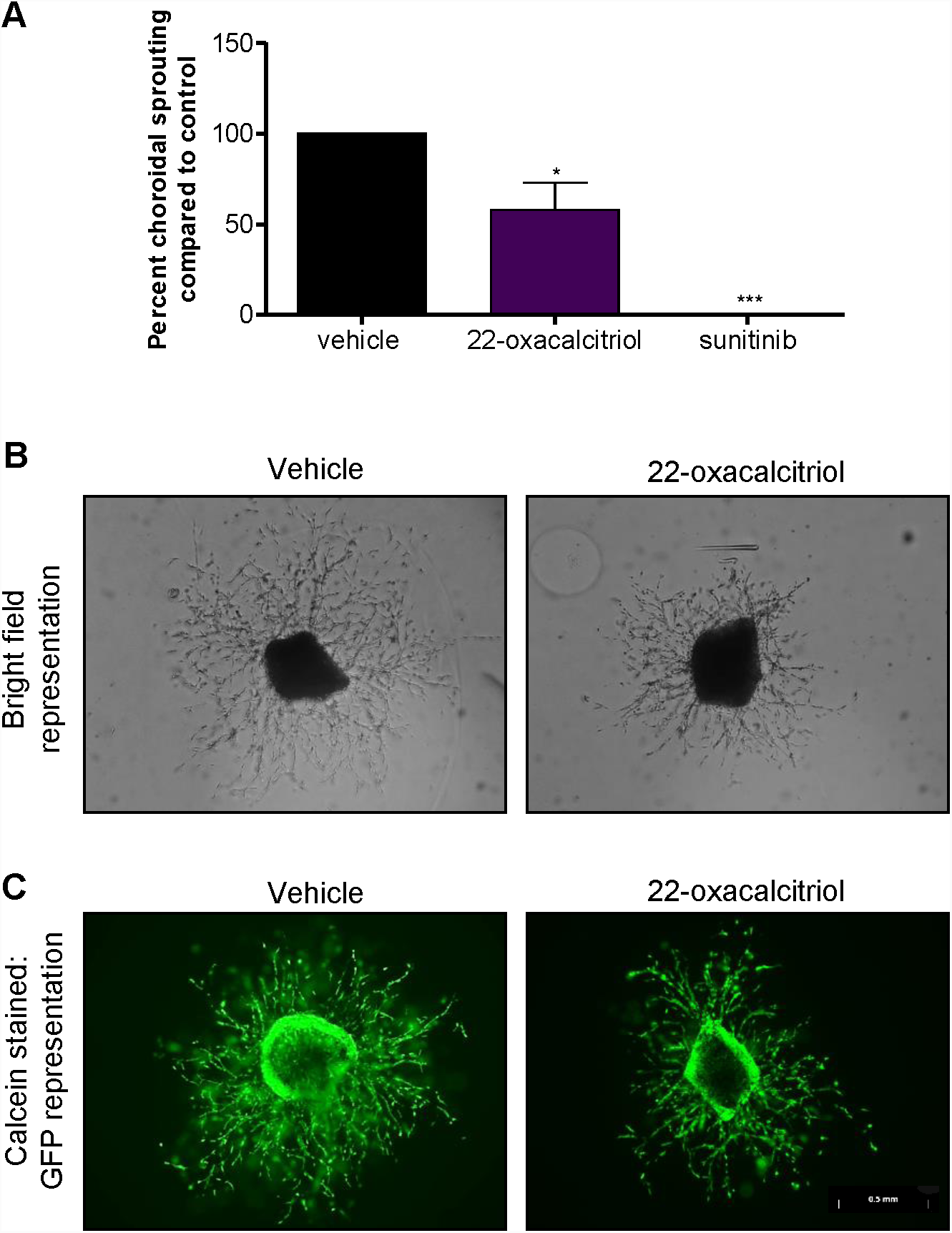
Vitamin D_*3*_ analogue attenuates mouse choroidal sprouting angiogenesis. Mouse RPE-choroidal fragments were cultured in Matrigel^®^ for 24 h and further cultured with 22-oxacalcitriol for 6 days. On day 7, samples were fixed and sprouting area quantified from phase contrast images using ImageJ freehand tool. **(A)** 22-oxacalcitriol and positive control sunitinib significantly attenuated choroidal sprouting angiogenesis. Graph showing mean percent sprouting area compared to vehicle control ±SEM; statistical analyses were performed using one-way ANOVA with Dunnett’s post hoc test; asterisk indicates *p≤0.05 and ***p≤0.001 and n =3 with up to 6 replicates per individual experiment. **(B)** Representative brightfield images of mouse choroidal sprouting after 7 days with indicated treatments. **(C)** Representative calcein stained images of mouse choroidal sprouting after 7 days with indicated treatments. Calcein staining of RPE-choroidal cultures confirms explant viability in vehicle and 22-oxacalcitriol treated explants. Scale bar represents 0.5 mm.

### Non-calcemic Vitamin D_3_ analogue 22-oxacalcitriol inhibits zebrafish choriocapillaris development

In zebrafish, the choriocapillaris starts developing by 24 hpf and becomes a perfused vascular network by 72 hpf. We treated zebrafish with ascending concentrations of 22-oxacalcitriol, between 24 and 72 hpf, and evaluated effects on choriocapillaris development. Using the number of interstitial pillars (ISPs) as a measure of angiogenic activity, the development of the choriocapillaris was markedly inhibited by 22-oxacalcitriol in a concentration-dependent manner. While 0.1 µM did not lead to reduced ISP formation compared to controls, treatment with 1 or 10 µM 22-oxacalcitriol led to a moderate reduction to 80.1% (±4.6) and 79.5% (±3.2) respectively of the ISPs found in control DMSO-treated eyes **(Figure 4B-C).** Importantly, treatment with 22-oxacalcitriol distorted the characteristic lobular vascular pattern of the choriocapillaris **(Figure 4C).** These findings clearly indicate that 22-oxacalcitriol inhibits choriocapillaris development in zebrafish.

**Figure 4:**
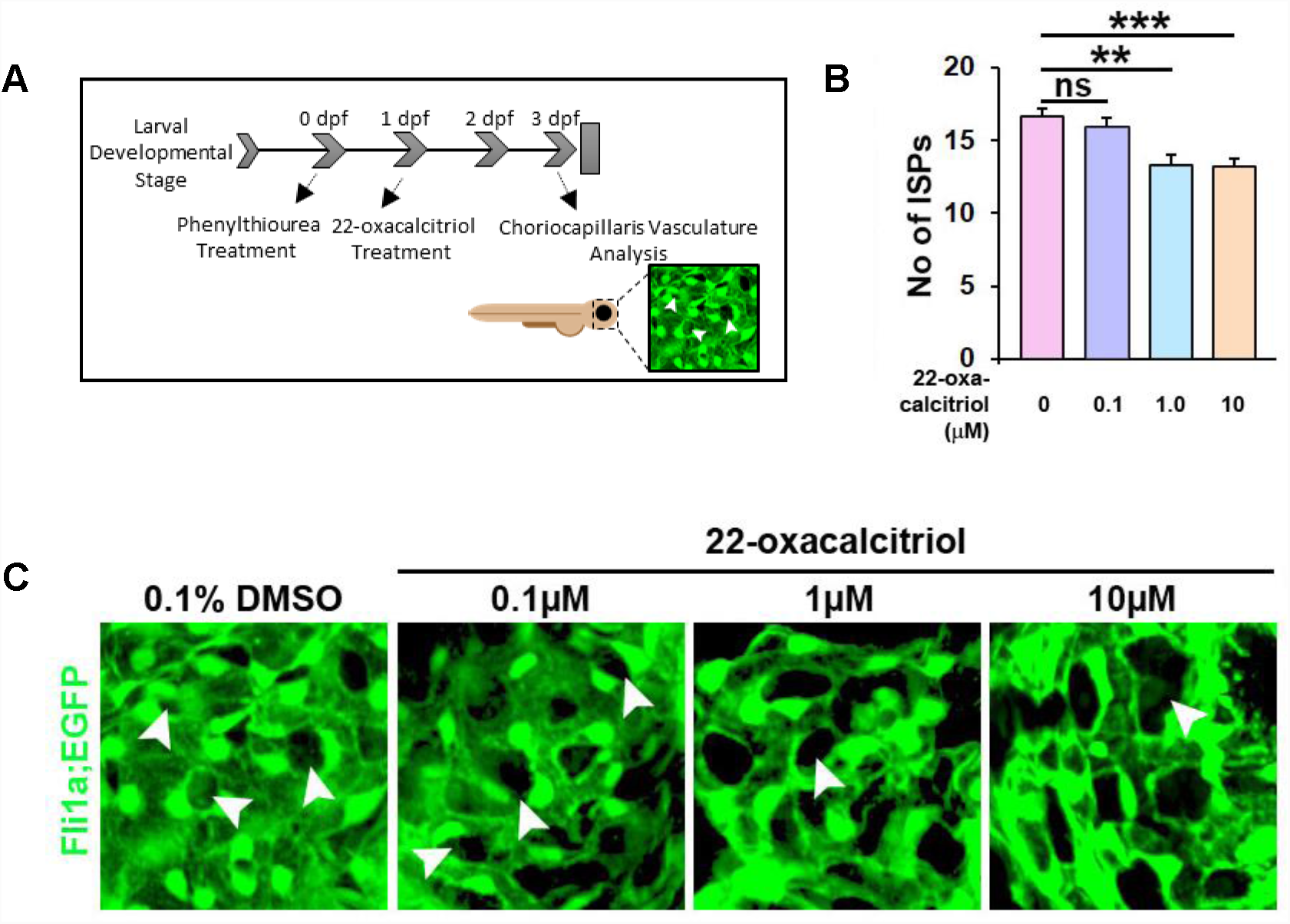
22-oxacalcitriol attenuates choriocapillaris development in zebrafish larvae. **(A)** Tg(Fli1a:EGFP)^y1^ transgenic zebrafish larvae were treated with vehicle control or 22-oxacalcitriol between 24-72 hpf. **(B)** Graph showing number of interstitial pillars (ISPs) as a measure of the extent of active vascular growth. Statistical analyses were performed using T-test; asterisk indicates ** p≤0.01 and *** p≤0.001 and n=2, N=20. ns, non-significant **(C)** Representative GFP images of choriocapillaris development in zebrafish larvae treated with vehicle control or 0.1-10 µM 22-oxacalcitriol treatment. Arrowheads indicate ISPs.

### Subcutaneous calcitriol treatment does not adversely affect adult mouse retinal structures or superficial retinal vasculature development

Prior to *in vivo* assessment of the anti-angiogenic activity of calcitriol, ocular safety was evaluated in adult C57BL/6J mice. Adult mice received a single 50 ng subcutaneous calcitriol or vehicle control treatment, and retinal histology was investigated 7 days later. Vehicle controls and calcitriol treated mice present with animal welfare scores comparable to uninjected mice, with mouse weight recorded daily **(Figure 5B)**. On day 7, mice were euthanised, eyes enucleated, fixed and sectioned. Ultra-thin toluidine blue stained cross sections were investigated for the presence of pyknotic nuclei and deviations in the highly-organised cell lamination of the eye. Calcitriol treatments appeared well tolerated in mice, with no observable difference between vehicle and calcitriol treated retinal structures **(Figure 5A)**.

**Figure 5:**
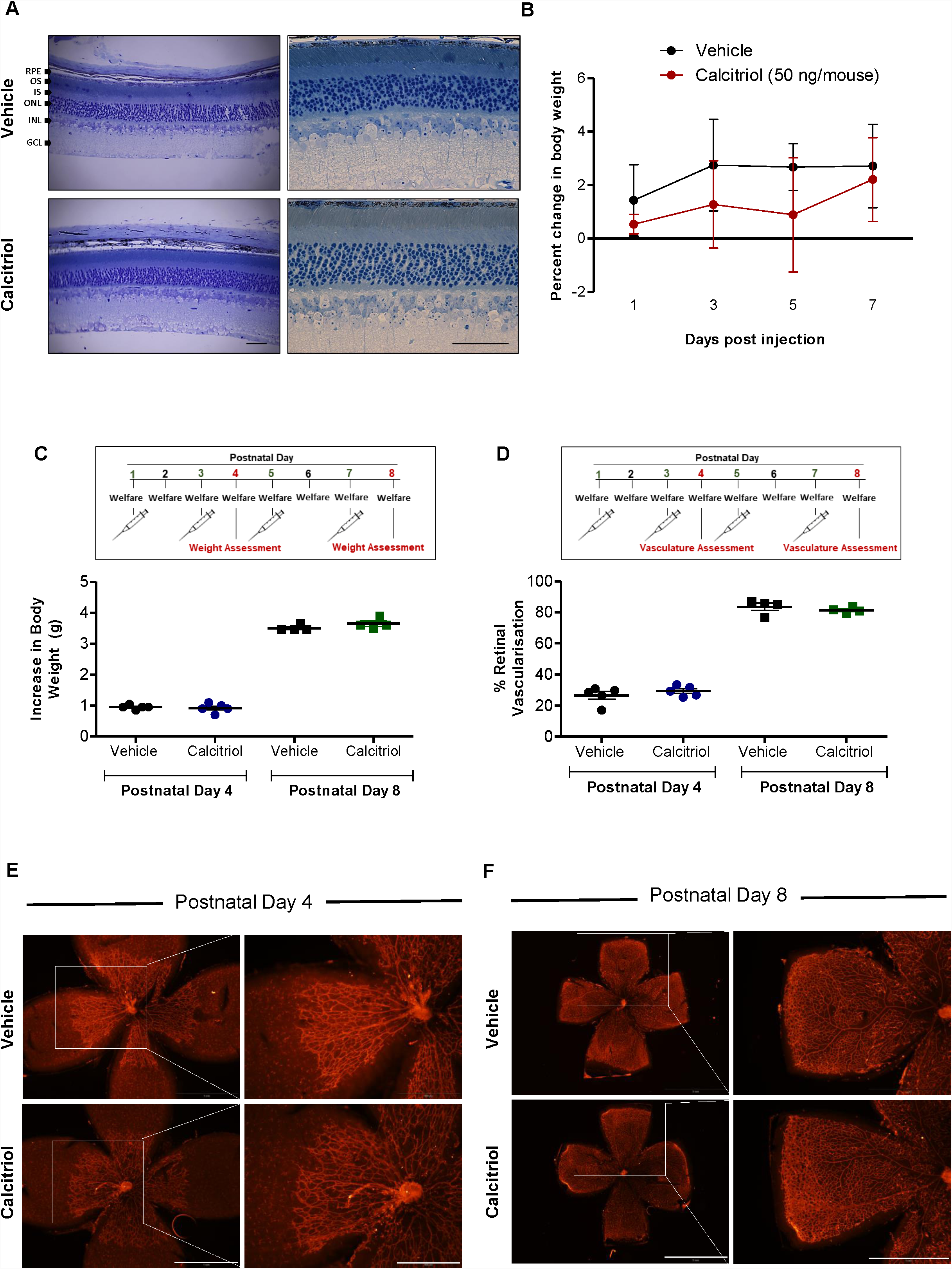
No ocular morphological defects were identified in mice treated with calcitriol and calcitriol treatment on alternating days does not attenuate superficial retinal vasculature development in C57BL/6J mice. **(A)** Toluidine blue stained ocular cross sections 7 days after vehicle or 50 ng calcitriol treatment show an absence of gross morphological defects and the presence of normal retinal lamination in ocular sections. RPE; retinal pigmented epithelium, OS; outer segment, IS; inner segment, ONL; outer nuclear layer, INL; inner nuclear layer, GCL; ganglion cell layer. Scale bar represents 0.5 mm. **(B)** Change in mouse body weight over 7 days was monitored as an indication of animal welfare. Graph shows change in body weight (g) ±SEM (n=3) in response to vehicle control or 50 ng calcitriol treatment. C57BL/6J mice at P1, P3, P5 and P7 received a s.c. injection of 3.75 ng calcitriol or vehicle control, welfare was monitored daily and superficial retinal vasculature development quantified at P4 or P8. **(C)** Change in mouse body weight between P1-4 or P1-8 was calculated, graph shows increase in body weight (g) from P1, ±SEM. No significant difference in body weight between 3.75 ng calcitriol and vehicle injected mice was identified at P4 or P8. **(D)** Superficial retinal vasculature area compared to retina area was calculated at P4 and P8. Scatter graph shows no significant difference between superficial retinal vasculature development at P4 or P8 between mouse pups treated with vehicle control or calcitriol. **(E-F)** Isolectin B4-Alexa Fluor 594 stained retinal flat mount image representations show no difference between superficial retinal vasculature development at P4 or P8 between mouse pups treated with vehicle control or 3.75 ng calcitriol. Scale bar represents 500 µm and 1 mm, left and right panels, respectively.

Retinal vasculature development after birth in mice provides a unique opportunity to study developmental angiogenesis. Normal mouse retinal vasculature growth is well documented, and drugs can inhibit this growth [34, 35]. Calcitriol attenuates zebrafish ocular developmental angiogenesis, therefore, we sought to investigate if this response translated to the mouse. To validate the model, the anti-angiogenic activity of positive control rapamycin was evaluated. C57BL/6J mouse pups were injected with 10 mg/kg subcutaneous rapamycin or vehicle control at P1, welfare monitored daily and superficial retinal vasculature development quantified at P4 **(Supp Figure 2)**. No significant difference in mouse weight gain was identified between rapamycin and vehicle treated animals at P1 or P4 **(Supp Figure 2A)**. Rapamycin treated mice presented with reduced superficial retinal vasculature area compared to vehicle control treated mice **(Supp Figure 2B)**.

Initial studies showed a single calcitriol injection was insufficient to induce an anti-angiogenic response in this model. C57BL/6J mice pups were injected with 3.75 ng subcutaneous calcitriol or vehicle control at P1, welfare monitored daily and superficial retinal vasculature development quantified **(Supp Figure 3)**. No significant difference in superficial retinal vasculature area between calcitriol and vehicle control administered mice was identified at P4 **(Supp Figure 3A & C)** or P8 **(Supp Figure 3B & C)**. A subsequent study performed subcutaneous calcitriol and vehicle control administration on P1, P3, P5 and P7. Mice pups presented with no welfare concerns and weight gain was comparable in vehicle control and calcitriol treated mice on P4 or P8 **(Figure 5C)**. Again, no significant difference in superficial retinal vasculature development between calcitriol and vehicle control treated mice was identified at P4 or P8 **(5D-F)**.

### 22-oxacalcitriol inhibits mouse laser-induced choroidal neovascularisation without adverse effects

Previously calcitriol was reported to reduce retinal neovascularization in the mouse oxygen induced retinopathy (OIR) model of retinopathy of prematurity [27]. To extend upon this finding, we evaluated the effect of calcitriol in L-CNV. To visualize L-CNV and consequential vascular leakage, *in vivo* optical coherence tomography (OCT) imaging and fluorescein angiography were performed. Choroidal neovascularisation was inhibited by 5 µg/kg/day calcitriol administered intraperitoneally **(Figure 6A-D)** and fluorescein angiography revealed vascular leakage of CNV lesions was reduced in calcitriol treated mice **(Figure 6B).** However, calcitriol treatment had adverse effects on body weight **(Figure 6D)**, perhaps due to hypercalcemia-induced toxicity [36].

**Figure 6:**
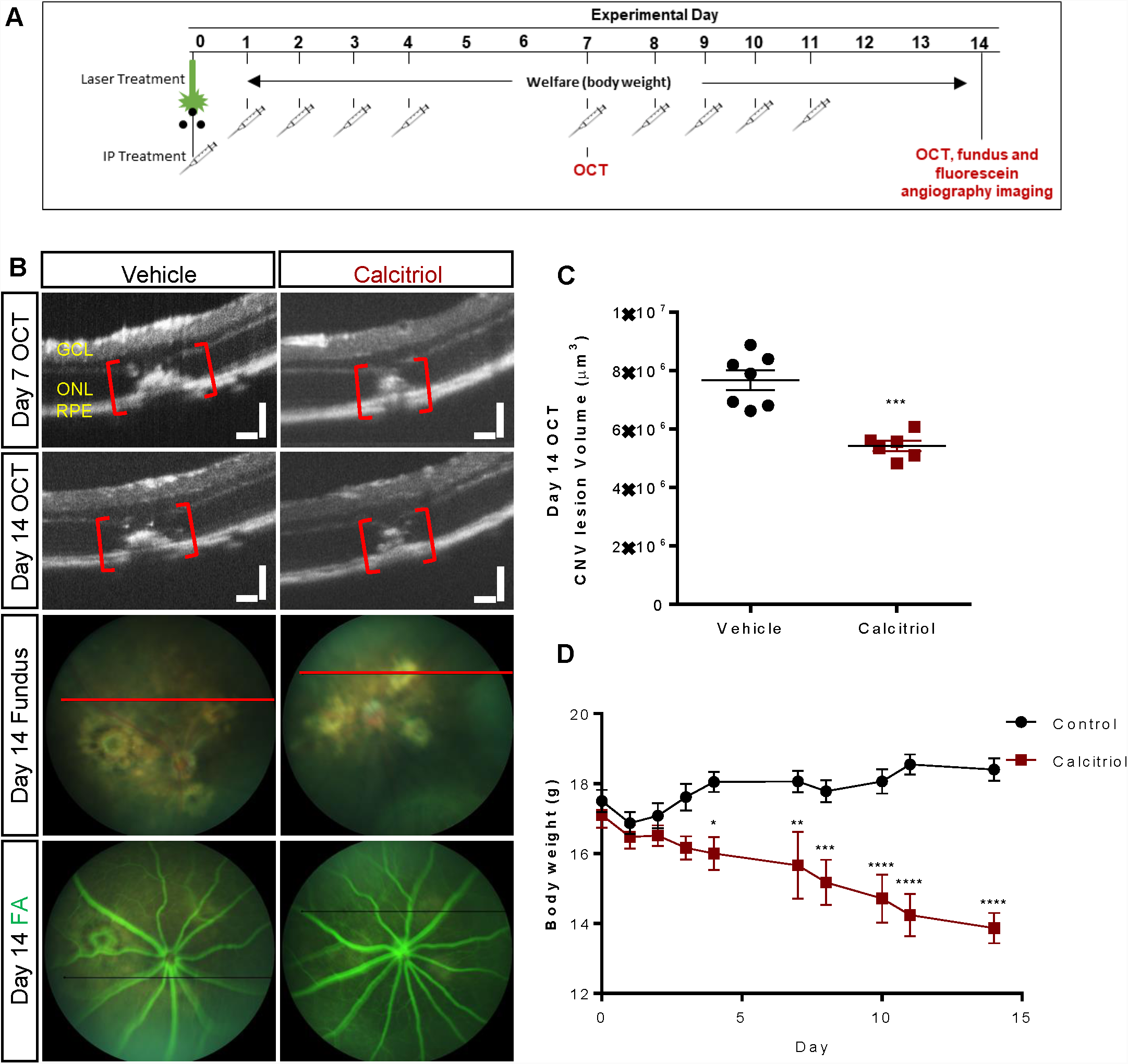
Effect of calcitriol on choroidal neovascularisation. **(A)** CNV was induced by laser on day 0 and calcitriol delivered i.p with a 5 days on/2 days off treatment regime. OCT, fundus and/or fluorescein angiography (FA) images were acquired on day 7 and/or 14. **(B)** Representative OCT, fundus and FA images on day 14. GCL: ganglion cell layer, ONL: outer nuclear layer, RPE: retinal pigment epithelium. Plane of OCT images shown as a red line on brightfield fundus images. Scale bars = 100 µm. **(C)** Quantification of the laser-induced CNV lesion volume calculated as an ellipsoid from OCT data, showing reduction in CNV lesions with calcitriol treatment on both day 7 and day 14. Calcitriol was delivered i.p. 5 µg/kg. Vehicle was almond oil. ***P=0.0002, Unpaired T-test (two tailed) with Welch’s correction, Mean ± SEM, n=6-7 eyes. **(D)** Effects of calcitriol treatment on body weight. *P<0.05, **P<0.01, ***P<0.001, ****P<0.0001, two-way repeated-measures ANOVA with Holm-Sidak post hoc tests. Mean ± SEM, n=6 animals.

Therefore, we sought to evaluate the effect of 22-oxacalcitriol, the non-calcemic bioactive analogue of calcitriol [37]. Notably, 22-oxacalcitriol (15 µg/kg/day) administered intraperitoneally inhibited vascular leakage **(Figure 7B)** and CNV lesion volume assessed based on OCT images **(Figure 7B-C)** without any adverse effect on body weight **(Figure 7D)**. CNV was also assessed immunohistochemically by staining vasculature using agglutinin and isolectin GS-IB4. CNV lesion volumes were reduced by 25-30% upon 22-oxacalcitriol treatment **(Figure 7F-G)**. QRT-PCR identified no significant difference between ocular Vegfa expression in mice treated with 22-oxacalcitriol or vehicle control **(Figure 7E)**.

**Figure 7:**
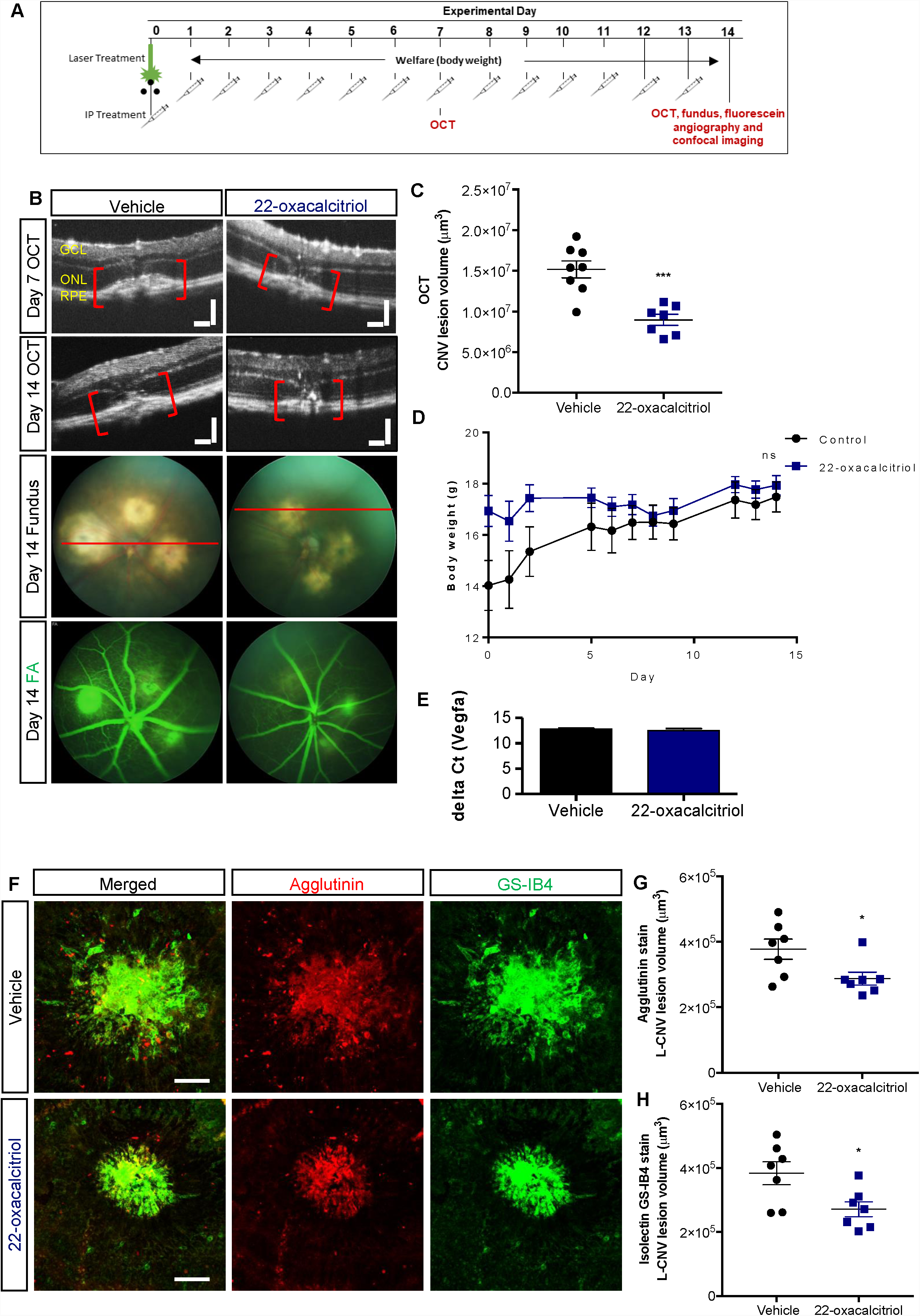
Effect of 22-oxacalcitriol on choroidal neovascularisation. **(A)** CNV was induced by laser on day 0 and 15 µg/kg 22-oxacalcitriol delivered i.p daily. Vehicle was 0.1% ethanol-PBS. OCT, fundus and/or fluorescein angiography (FA) images were acquired on day 7 and 14. **(B)** Representative OCT, fundus and FA images on day 14. GCL: ganglion cell layer, ONL: outer nuclear layer, RPE: retinal pigment epithelium. Plane of OCT images shown as a red line on brightfield fundus images. Scale bars = 100 µm. **(C)** Quantification of the laser-induced CNV lesion volume calculated as an ellipsoid from OCT data, showing reduction in CNV lesions with 22-oxacalcitriol treatment on both day 7 and day 14; day 14 data shown. ***P=0.0003, Unpaired t-test (two tailed) with Welch’s correction, Mean ± SEM, n=7-8 eyes. **(D)** Effects of 22-oxacalcitriol treatment on body weight. ns, non-significant, two-way repeated-measures ANOVA. Mean ± SEM, n=6 animals. **(E)** *Vegfa* mRNA expression in the mouse eye was quantified after daily 22-oxacalcitriol or vehicle control for 14 days. *Vegfa* expression was consistent in vehicle control and 22-oxacalcitriol treated eyes. **(F)** Representative images from confocal microscopy of agglutinin and GS-IB4 stained CNV lesions 14 days post L-CNV. Scale bars = 100 μm. **(G-H)** Quantification of CNV lesion volume using Z-stack confocal images. *P<0.03, Unpaired T-test (two tailed) with Welch’s correction, Mean ± SEM, n=7 eyes.[23].

In summary, calcitriol and 22-oxacalcitriol attenuate mouse *ex vivo* choroidal vasculature sprouting, 22-oxacalcitriol reduces choriocapillaris development in zebrafish larvae and both calcitriol and 22-oxacalcitriol inhibit choroidal neovascularisation in a mouse model with features of nAMD.

## DISCUSSION

The ocular vasculature systems support retinal development and visual function. Pathological ocular neovascularisation is a hallmark in numerous diseases causing vision loss, including nAMD. Anti-angiogenics have revolutionised nAMD treatment, yet their long-term safety profiles remain controversial and they require frequent intravitreal injection due to their molecular weights greater than 50 kDa [38, 39]. Here, we investigated the anti-angiogenic activity and safety of agonists targeting the VDR in zebrafish larvae, human cell cultures and murine models. We report four significant findings. First, we demonstrate the *ex-vivo* ocular anti-angiogenic activity of calcitriol and 22-oxacalcitriol in a physiological mouse model of angiogenesis. Second, 22-oxacalcitriol exerts ocular anti-angiogenic activity in a zebrafish model of choroidal vasculature development. Third, calcitriol and 22-oxacalcitriol attenuates neovascularisation in an *in-vivo* pathological mouse model with features of nAMD. Finally, we show 22-oxacalcitriol, a vitamin D agonist not associated with hypercalcemia, to present with an improved safety profile compared to calcitriol in human RPE cells and mice. Together this supports the potential of vitamin D agonists, particularly 22-oxacalcitriol, as anti-angiogenic agents for the treatment or prevention of ocular angiogenic disorders.

A combination of pathological changes to the photoreceptors, RPE, Bruch’s membrane and choroid underpin visual impairment in nAMD. Therefore, modelling such a complex disease *in vitro* can be challenging. The *ex vivo* choroidal sprouting angiogenesis model is multi-cellular, comprising RPE cells, endothelial cells, macrophages and pericytes [22]. The choriocapillaris is the vasculature located immediately posterior to the RPE and highly sensitive to VEGF-A signalling during development and in pathologies such as AMD [1]. In initial studies, calcitriol attenuated *ex vivo* choroidal sprouting angiogenesis. This anti-angiogenic response is consistent with previous studies wherein calcitriol inhibits angiogenesis in a chick chorioallantoic membrane (CAM) model [40], an OIR model [27] and a transgenic mouse model of retinoblastoma [40]. Interestingly, here calcitriol did not inhibit *in vitro* tubule formation in human dermal- or retinal-derived endothelial cells. This is contrary to previous findings which demonstrated calcitriol to inhibit mouse retinal endothelial cell capillary network formation on a basement membrane [27]. Interestingly, *Bao et al* reported calcitriol to exert no effect on HUVEC tubule formation, yet attenuated HUVEC tubule formation when stimulated with prostate cancer cell conditioned medium [41]. These differences suggest that the anti-angiogenic activity of calcitriol is context- or environmental-dependent. As the choroidal explant assay employed here is multi-cellular in nature, it is plausible that anti-angiogenic responses are not directly induced at the endothelial cells, but instead through regulation from neighbouring cells.

Calcitriol treatment induces oedema and impairs visual function in zebrafish larvae [42]. Therefore, the safety profile of calcitriol was assessed here in a mammalian system. In mice, calcitriol treatment was well-tolerated in developing pups with no ocular morphological welfare concerns or weight loss at selected dose. We hypothesised that *in vivo* calcitriol would stall development of the mouse retinal vasculature, correlating with previous observations in zebrafish larvae [42]. However, mouse retinal vascular plexus development at P4 or P8 was not attenuated by 0.00375 μg calcitriol subcutaneously administered as a single dose or repeatedly on alternating days. The lack of efficacy could be a consequence of suboptimal dosing or short drug half-life (t_1/2_). Indeed, the calcitriol t_1/2_ is only a few hours, ∼100 times quicker than 25(OH)D_3_ [43]. Alternatively, lack of anti-angiogenic activity could result from poor distribution to the eye, suboptimal vehicle selection or the early developmental status of the retinal vasculature. Calcitriol *in vivo* was previously reported to inhibit retinal neovascularization in a pathological OIR model of retinopathy of prematurity at a concentration of 5 μg/kg via intraperitoneal administration daily from P12 to P17 [27]. Subsequently, we hypothesised that calcitriol may exert an anti-angiogenic effect *in vivo* on the choroidal vasculature, consistent with our data in the *ex vivo* choroidal explants. Significantly, 5 µg/kg calcitriol administered intraperitoneally daily attenuated L-CNV in adult mice. However, the short calcitriol t_1/2_ suggests that frequent administration is required for chronic conditions [43]. Of further concern, the calcitriol-treated arm in the L-CNV study presented with reduced body weight, a response also reported by Albert *et al* in OIR studies [27]. Weight loss in calcitriol treated animals is likely a consequence of hypercalcemia.

Vitamin D regulates calcium mobilisation, therefore high-dose vitamin D treatment can induce hypercalcemia. VDR agonists including 22-oxacalcitriol were developed to reduce calcemic responses. 22-oxacalcitriol is a calcitriol analogue with an oxygen substituted for carbon at position 22, and is approved for the treatment of psoriasis due to its therapeutic activity and reduced calcaemic responses [44]. Interestingly, 22-oxacalcitriol inhibits CAM angiogenesis in a dose-dependent manner [45]. Here calcitriol decreased ARPE-19 cell viability, a result not observed with 22-oxacalcitriol. Differing responses induced by vitamin D and analogues are the result of altered protein binding, metabolism, receptor affinity, dimerization and co-regulator recruitment [44]. 22-oxacalcitriol is reported to have reduced vitamin D-binding protein affinity, up to 500 times lower calcitriol [46].

22-oxacalcitriol attenuates zebrafish hyaloid vasculature development, a lens associated system which metamorphs into a retina-associated vasculature system [42, 47]. Here, 22-oxacalcitriol attenuated both *ex vivo* mouse microvascular sprouting in choroidal explants and choroidal vasculature development in zebrafish larvae. Thus, our revised hypothesis was that 22-oxacalcitriol can exert an anti-angiogenic effect on the choroidal vasculature *in vivo* without adverse effects linked to hypercalcemia. Here, our novel findings demonstrate 22-oxacalcitriol to exert significant anti-angiogenic activity in the mouse L-CNV model. Importantly, weight loss did not occur in the 22-oxacalcitriol treatment arms.

Vitamin D traditionally mediates its effects though the VDR, a nuclear receptor expressed diversely throughout the body which regulates the transcription of hundreds of genes [44]. The anti-angiogenic effects of vitamin D appear VDR-dependent. First, knockout of the VDR affects tumour vasculature integrity, resulting in vessel enlargement and reduced pericyte coverage [48]. Second, increased expression of pro-angiogenic factors including VEGF has been identified in tumours from VDR KO mice [48]. Third, calcitriol attenuates oxygen-induced neovascularisation in mice, a response reduced in VDR KO mice [49]. We previously reported calcitriol treatments to regulate VEGF expression in the developing eye [42]. However, here in the L-CNV model, ocular VEGF_A_ expression was not altered by 22-oxacalcitriol. This is in line with Albert *et al* who found VEGF protein expression to be comparable in calcitriol and vehicle control mice eyes during the OIR model of ocular neovascularisation [27]. Angiogenesis is driven by an array of factors. Indeed, vitamin D regulates a plethora of other angiogenic factors including HIF1α, IL-8, TGF-β, bone morphogenetic protein-2A, endothelin 1, cysteine-rich angiogenic inducer 6, midkine, MMP-2 and MMP-9 [41, 50, 51]. Thus, additional studies are needed to elucidate the anti-angiogenic mechanism and VEGF dependency of responses induced by vitamin D in an ocular setting.

In summary, we present original findings on the anti-angiogenic efficacy of VDR agonists in the eye using an evolutionarily diverse range of *in vivo* and *ex vivo* models. Notably, in a mouse model of pathological choroidal neovascularisation, 22-oxacalcitriol is well-tolerated and significantly reduces the size of vascular lesions. This drug represents a promising therapeutic or preventative treatment for ocular angiogenic disorders associated with choroidal vasculopathies.

## Acknowledgments: Funding Statement

### This work was supported by

Health Research Board, HRB-POR-2013-390, Irish Research Council, GOIPG/2015/2061, NIH/NEI R01EY025641, Retina Research Foundation, The foundation Jeanssons Stiftelser, The Magnus Bergvall foundation, andThe Swedish Eye foundation

### Conflict of Interest Statement

No conflicting interest.

### Author Contribution statement

Merrigan SL performed and analysed mouse choroidal sprouting angiogenesis assays, in vitro assays and mouse models of retinal vasculature development. Ali Z and Jensen LD performed and analysed choriocapillaris development assays in zebrafish. Park B and Corson TW performed and analysed mouse models of laser-induced choroidal neovascularisation. Supervision, resources, experimental designs and result interpretations were carried out by Kennedy BN. Kennedy BN and Merrigan SL drafted manuscript with input from Park B, Ali Z, Jensen LD and Corson TW.

### Ethical statement and approvals

Approval for the use of post-mortem mouse tissue for research purposes was granted by the UCD Animal Research Ethics Committee (AREC) committee, AREC-16-31-Kennedy. Mouse developmental and safety studies were conducted with approval by the UCD AREC committee (AREC-15-07-Kennedy) and the Health Products Regulatory Authority (AE18982/P064). Mouse L-CNV experiments followed the guidelines of the Association for Research in Vision and Ophthalmology Statement for the Use of Animals in Ophthalmic and Visual Research and were approved by the Indiana University School of Medicine Institutional Animal Care and Use Committee. Zebrafish choroidal angiogenesis studies were approved by the Linköping Animal Research Ethics Committee (N89/15). All experiments were performed in accordance with relevant guidelines and regulations.

