## Supplementary figures and images for "Calcitriol and Non-Calcemic Vitamin D Analogue, 22-Oxacalcitriol, Attenuate Developmental and Pathological Ocular Angiogenesis Ex Vivo and In Vivo"

### Supplementary Figures

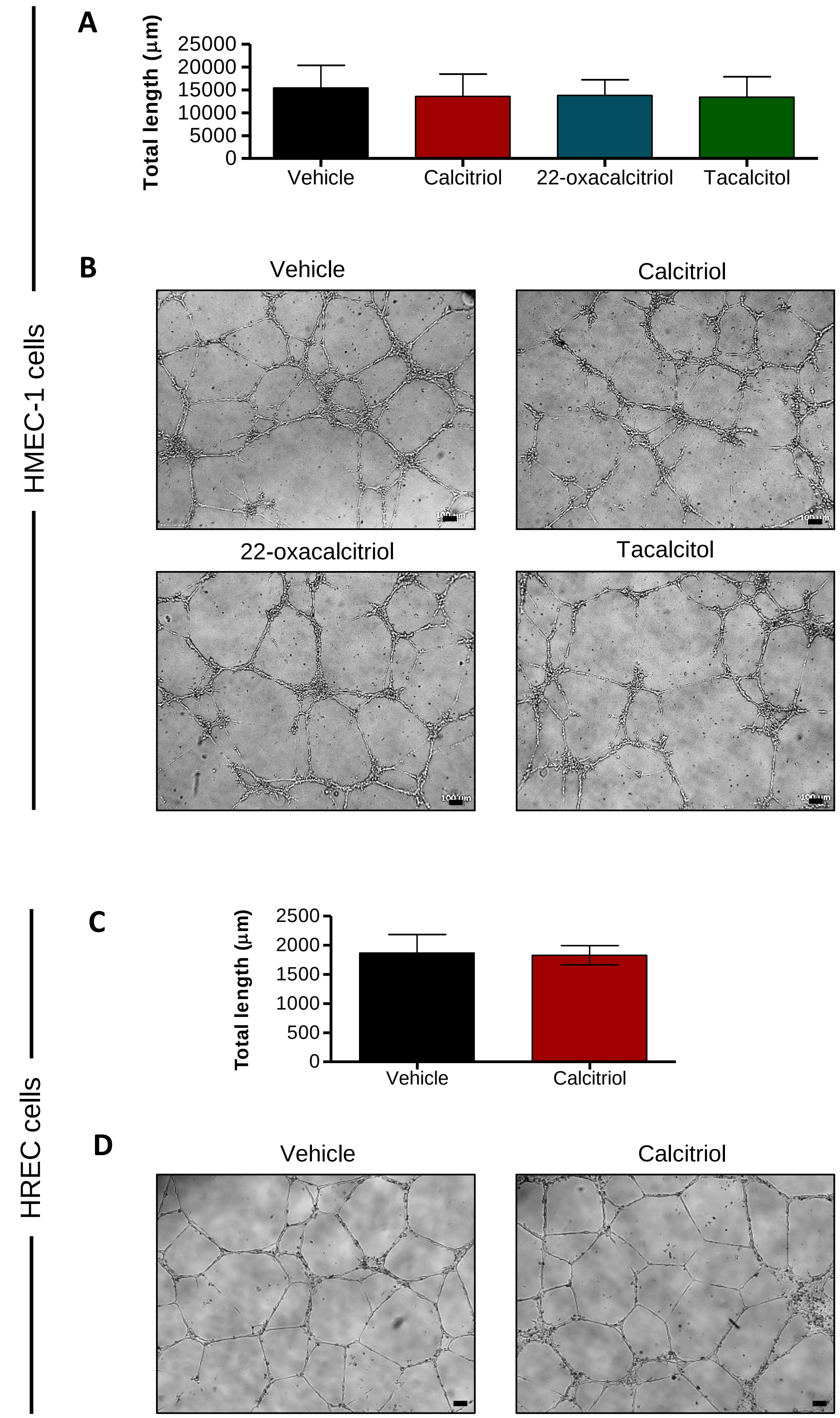

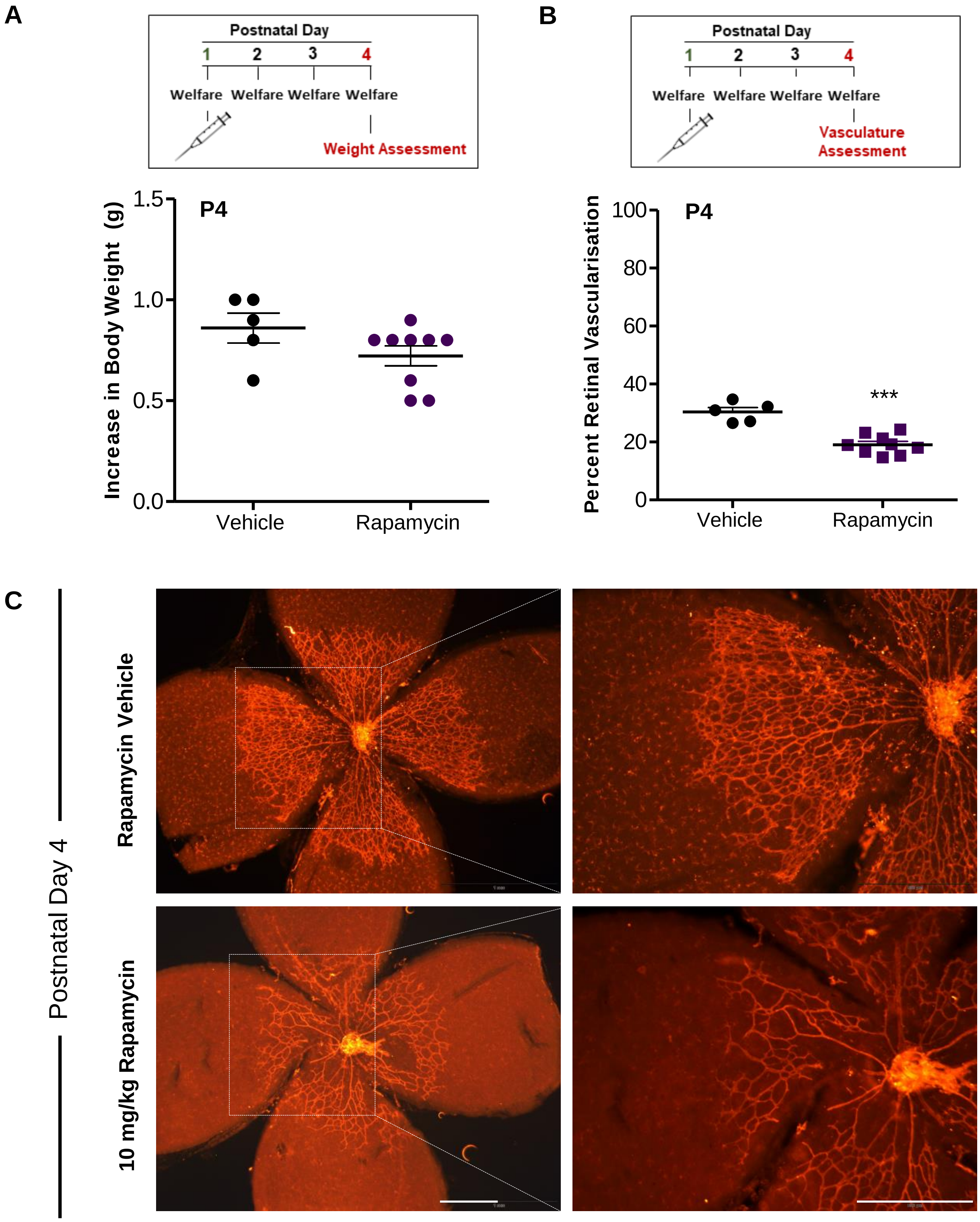

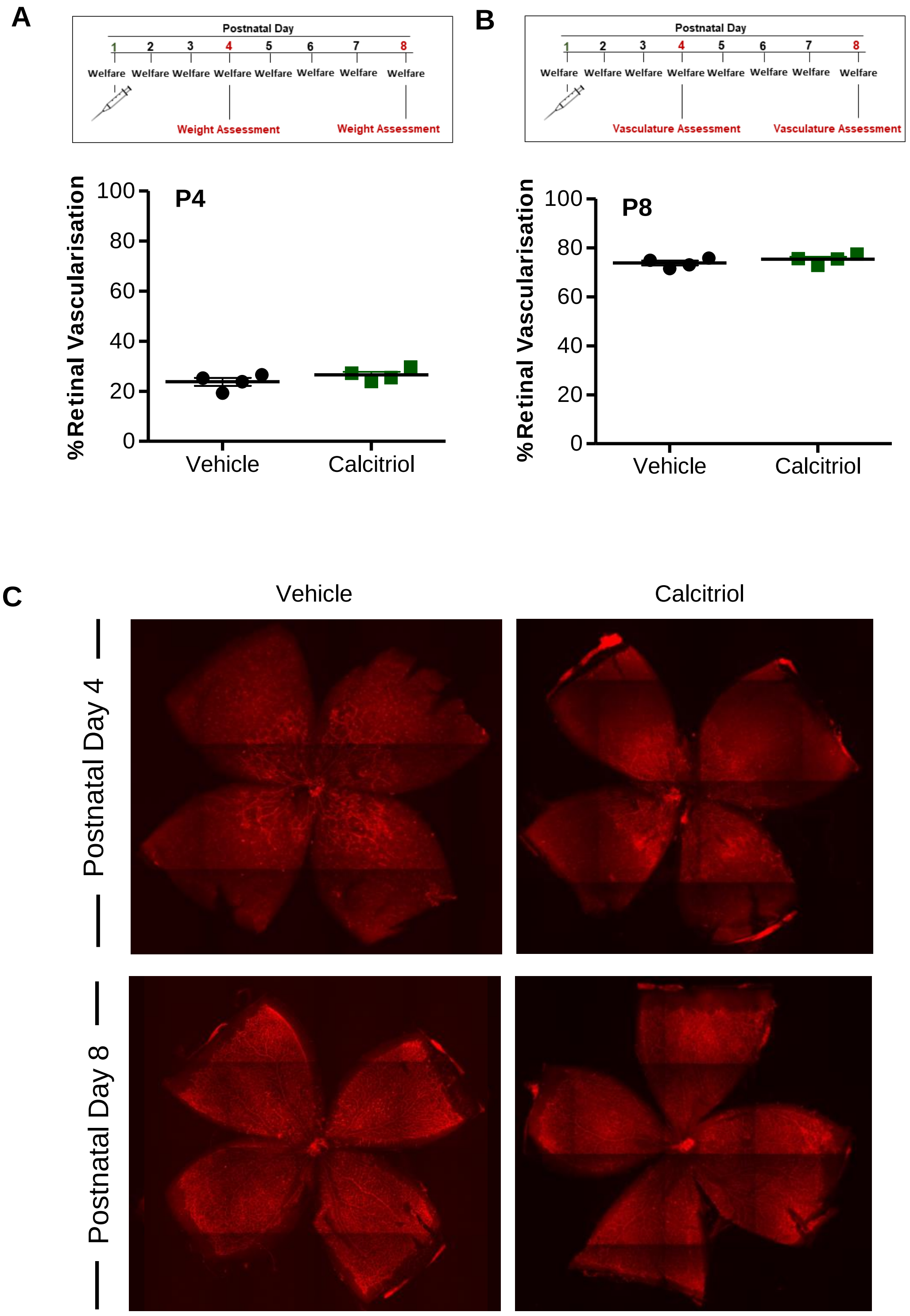
