## Supplementary Text for "Calcitriol and Non-Calcemic Vitamin D Analogue, 22-Oxacalcitriol, Attenuate Developmental and Pathological Ocular Angiogenesis Ex Vivo and In Vivo"

### Methods

#### *In vitro tubule formation assay*

*Vitamin D receptor agonist dosing:* 10 mM drug stocks in DMSO were prepared. Working solutions to 100 µM were prepared in sterile filtered $dH_2O$ (Fisher scientific, EMD Millipore Millex-GP Sterile Syringe Filters) and final dilutions to treatment concentrations made in cell type dependent medium.

*Cell maintenance:* Dermal derived HMEC-1 cells were cultured in MCDB-131 Medium supplemented with 10% fetal bovine serum (Gibco), 2 mM L-glutamine (Gibco), 100 units/mL penicillin-streptomycin (Gibco), 1 µg/mL hydrocortisone (Sigma) and 10 ng/mL epidermal growth factor (BD Biosciences). HRECs (ACBR 181, Cell Systems) were cultured in CSC Complete Medium (Cell Systems) supplemented with Bac-Off® (antibiotic) and CultureBoost™ (animal derived growth factors). HRECs were cultured as per supplier specifications.

*In vitro* tubule formation assay was performed as per manufacturer's instructions (ibidi, µ-Slide Angiogenesis). Ibidi µ-slide angiogenesis plates were coated with 10 µl Matrigel® (BD Biosciences) and incubated at 37°C (5% $CO_2$, 95% $O_2$) for 30 min. Each Matrigel® coated well was seeded with 7.5 $x10^3$ HREC or HMEC-1 cells in cell dependent medium (25 µl). Drug treatments were prepared in cell type-dependent medium with a final volume of 25 µl, and concentrations as indicated. Plates were incubated at 37°C (5% $CO_2$, 95% $O_2$) for 16 h.

A representative image for each well was acquired by phase contrast microscopy (Zeiss Axiovert 200 M microscope). Tubule length was measured manually using Zeiss Axiovision image analysis software or Angiogenesis Analyzer for ImageJ employed for unbiased measurements. Angiogenesis Analyzer generates 22 measurements of tubule properties from RGB coded images, here total length was selected for analyses and average ±SEM plotted.

#### *Mouse model of retinal vasculature development*

*Rapamycin Dosing:* Positive control rapamycin was prepared as previously reported [1]. Rapamycin was dissolved to 25 mg/ml in DMSO and working dilutions prepared to 1 mg/ml in 4% EtOH, 5% Tween 80 and 5% Kollisolv(R) PEG E 400 (Sigma-Aldrich). Pups received a single 10 mg/kg rapamycin or vehicle control s.c. treatment on P1. Animal welfare was monitored daily until P4 when mouse pups were euthanised by cervical dislocation, eyes immediately enucleated and fixed with 4% PFA overnight at 4°C.

*Calcitriol Dosing:* Calcitriol single injection study: Pups received a single 3.75 ng calcitriol or vehicle control s.c. treatment on P1. Animal welfare was monitored daily until P4 (50% of pups) or P8 (50% of pups). On P4 and P8 mouse pups were euthanised by cervical dislocation, eyes immediately enucleated and fixed with 4% PFA overnight at 4°C.

### Figure Legends

***Supp figure 1: HMEC-1 and HREC tubule formation properties are unchanged by VDR agonists.*** **(A-B)** HMEC-1 cells were seeded on a Matrigel® coated µ-slide with 10 µM 22-oxacalcitriol, calcitriol, tacalcitol, or 0.1 % DMSO vehicle control for 16 h. Phase contrast images were acquired, and Angiogenesis Analyzer for ImageJ utilised to determine total tubule length in the analysed area. **(A)** Graph showing mean ±SEM. No significant differences in tubule formation properties were identified compared to vehicle control, n=3 with N=2/n. **(B)** Representative images of tubule formation on a Matrigel® coated µ-slide with 16 h vehicle control or VDR agonist treatments. Scale bar represents 100 µm. **(C-D)** HREC cells were seeded on a Matrigel® coated µ-slide with 10 µM calcitriol or vehicle control for 16 h. Phase contrast images were acquired, and Angiogenesis Analyzer for ImageJ utilised to determine sum of the length of the branches in the analysed area. **(C)** Graph shows mean compared to vehicle control ±SEM. No significant differences in tubule formation properties were identified, n=2 with N=2/n. **(D)** Representative images of HREC tubule formation on a Matrigel® coated µ-slide with 16 h calcitriol or vehicle control treatment. Scale bar represents 100 µm.

***Supp figure 2: mTOR inhibitor, rapamycin, attenuates superficial retinal vasculature development in C57BL/6J mice.*** C57BL/6J mice at P1 received a single subcutaneous injection of 10 mg/kg rapamycin, welfare was monitored daily and superficial retinal vasculature development quantified at P4. **(A)** Gain in mouse body weight between P1 and P4 was calculated as an indication of animal welfare. Graph shows increase in body weight (g) ±SEM, no significant difference in body weight between rapamycin and vehicle injected mice was identified. **(B)** Superficial retinal vasculature area compared to retina area was calculated at P4, rapamycin significantly reduced retinal superficial vasculature development. Scatter plot shows percent vasculature coverage compared to total retinal area ±SEM; unpaired t-test statistical analyses, asterisks signify ***$p \leq 0.001$; n=5-9. **(C)** Images of retina flat-mounts, isolectin B4-Alexa Fluor 594 stained, showing superficial retinal vasculature in mouse pups at P4. A clear reduction in superficial retinal vasculature can be seen in rapamycin compared to vehicle control treated mice. Scale bar represents 500 µm.

***Supp figure 3: A single calcitriol treatment does not attenuate superficial retinal vasculature development in C57BL/6J mice.*** C57BL/6J mice at P1 received a single subcutaneous injection of calcitriol or vehicle, welfare was monitored daily and superficial retinal vasculature development quantified at P4 or P8. **(A-B)** Superficial retinal vasculature area compared to retina area was calculated at P4 and P8. Scatter graph shows no significant difference between superficial retinal vasculature development at **(A)** P4 or **(B)** P8 in mouse pups treated with vehicle control or 3.75 ng calcitriol. **(C)** Image representations of retina flat-mounts isolectin B4-Alexa Fluor 594 stained, showing superficial retinal vasculature in mouse pups at P4 and P8 after vehicle or 3.75 ng calcitriol treatment. No significant reduction in superficial retinal vasculature was observed following calcitriol treatment.
